# Modeling Epidemics: A Primer and Numerus Software Implementation

**DOI:** 10.1101/191601

**Authors:** Wayne M. Getz, Richard Salter, Oliver Muellerklein, Hyun S. Yoon, Krti Tallam

**Author notes:** Correspond Author Wayne M Getz.

## Abstract

Epidemiological models are dominated by SEIR (Susceptible, Exposed, Infected and Removed) dynamical systems formulations and their elaborations. These formulations can be continuous or discrete, deterministic or stochastic, or spatially homogeneous or heterogeneous, the latter often embracing a network formulation. Here we review the continuous and discrete deterministic and discrete stochastic formulations of the SEIR dynamical systems models, and we outline how they can be easily and rapidly constructed using the Numerus Model Builder, a graphically-driven coding platform. We also demonstrate how to extend these models to a metapopulation setting using both the Numerus Model Builder network and geographical mapping tools.

## 1 Introduction

In a recent comprehensive review of epidemiological models, Smith et al. [1] trace the development of systems of differential equations used over the past 100 years to study disease processes. Once the purview of mathematicians, physicist and engineers, dynamical system formulations of epidemic processes are increasingly being used by epidemiologists, ecologists and social scientists to study the potential for disease pandemics to threaten the lives of humans, domesticated animals and plants, and all organisms across the globe.

Underpinning all dynamical systems models of epidemiological outbreaks and endemics disease are formulations based on the concept of an SEIR progression (Figure 1), whereby susceptible individuals in disease class S enter disease class E on exposure to a pathogen (i.e., infected but not yet infectious themselves). Individuals in class E then transfer, after a period of latency, into the class of infectious individuals, I, only to transfer on recovery or death to a removed class R. The recovered individuals may have some temporary, though possibly long term, immunity (V) and death (D) can arise from both natural and disease-induced causes (i.e., R=V+D). Variations and elaboration of the the SEIR process may include birth and death processes, other demographic class such as age [2] and sex [3], spatial structure [4, 5], and the genetic structure of hosts and pathogens [6,7]. Further, stochastic formulations of SEIR models [8, 9] are increasingly being used to explore the inherent and very important stochastic aspects of epidemics including, probabilities of fadeouts versus breakouts in the early stages of epidemics, and logistical planning necessary for managing epidemics of uncertain ultimate size.

**Figure 1:**
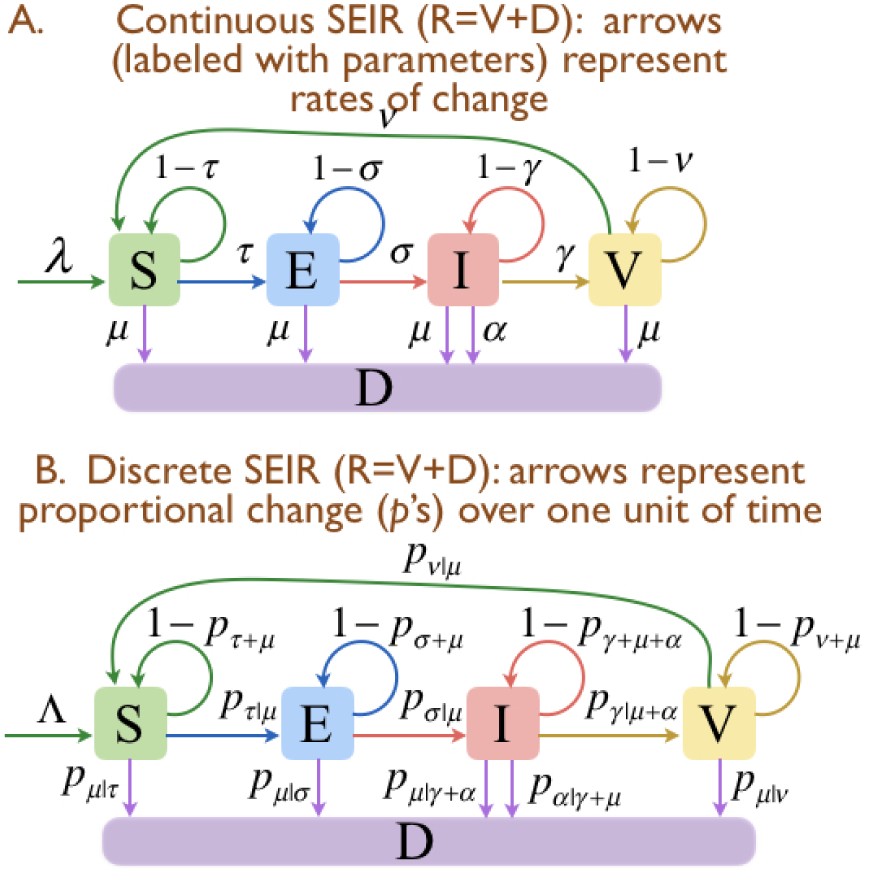
Flow diagrams for the basic SEIV continuous (A) and discrete (B) time models with transition rates *τ, σ, γ* and *ν* from disease classes S to E, E to I, I to V and V back to I, respectively. The class of dead individuals D together with immune individuals V make up the historically defined “removed class” R. In the continuous-time formulation, λ is the rate at which new individuals are recruited to the susceptible population (births or emigration), *μ* and *α* are natural and disease-induced mortality rates respectively. In the discrete-time formulation, a competing rates approach is used to derive transition proportions *p*, with identifying subscripts, as described in the text.

Hethcote [10] provided a comprehensive review of the fundamental deterministic dynamics of SEIR models and their elaboration to include an M class (that is, individuals born with maternally provided immunity that wears off over time), and elementary characterizations of birth and death process. His results have been considerably extended to elaborated SEIR models that include more complex characterizations of births and of deaths, and some age-structure - with a strong focus in the mathematics literature on the existence and stability of outbreak and endemic equilibria [11]. Analyses of stochastic SEIR models and elaborations remain more challenging, though some analytical results do exist. Most stochastic analyses, however, are computationally intensive and the results numerical rather than analytical. In addition, unlike deterministic SEIR models which are readily fitted to data, it remains a challenge to fit stochastic models to data, with new approaches involving concepts well beyond first courses in calculus or linear algebra.

Given that an SEIR structure underpins all epidemiological models, whether deterministic, stochastic, or even agent-based, a succinct, pedagogical review of SEIR modeling is useful. In particular a clear exposition of SEIR models for the non-mathematician—by which we mean, scientists who have some understanding of calculus, but do not have formal training in dynamical systems theory, or the numerical techniques to competently build and implement computational models. In addition, the literature lacks expository articles that elucidate for biologists and social scientists the relationship among continuous and discrete SEIR models (but see [12]), their stochastic elaborations in systems and agent-based (i.e., individual-based) computational settings [13,14], as well as extensions to metapopulation settings [15]. These are lacunae we hope to fill with this paper, while at the same time providing those scientists who are looking for fast, reliable ways to obtain and modify code needed to address their epidemiological models with a means to do so in the context of the Numerus Model Builder software development platform.

Finally, given the centrality of dynamic epidemic models to containment of outbreaks, policy formulation and response logistics, modeling tools are needed that can be used by healthcare professionals not trained in computational methods to carry out containment policy, and response analyses. Thus, our strong focus is on how to use the Numerus Model Builder and demonstrate its use to address these issues in the context of epidemics that are spatially structured, such as the recent outbreak of Ebola in West Africa [16,17].

## 2 Homogeneous SEIR Formulations

### 2.1 Continuous Deterministic Models

SEIR infectious disease models are based on dividing an otherwise homogeneous population into the following disease classes: susceptible (S), exposed (E; infected but not yet infectious), infectious (I), and Removed (R) individuals, the latter comprising either dead (D) or recovered with immunity (V; for vaccinated, though naturally so) that may wane over time. Through-out, we use the roman font S, E, I, V and D to name the class itself and the italic font *S*, *E*, *I*, *V* and *D* to refer to the variables representing the number of individuals in the corresponding classes. The assumption of homogeneity implies that age and sex structure are ignored. We incorporate population spatial structure—as would be found in countries comprising of a network of cities, towns, and villages—into a metapopulation framework [15,18], if we assume that a set of homogeneous subpopulations can be organized into a network of subpopulations, among which individuals move in a fashion that reflects appropriate movement rates (e.g., propensity to move as a function of age and sex [19])and geographical factors (e.g., distances, geographical barriers,
desirability of possible destinations).

If the time scales of the epidemic and movement processes among subpopulations, including disease induced mortality, are much faster than the time scale of the background population demography (births, recruitment, natural mortality and population level migration) then we can ignore the demography; otherwise we cannot. For example, in the case of influenza, epidemiological and local movement processes involve noticeable changes at the scale of weeks, while demographic changes in the underlying population itself (beyond epidemic disease induced death rates) are obvious only at the scale of years. In this case, we can ignore natural births and deaths, and focus on epidemic processes alone.

In the context of an epidemic occurring in a single homogeneous population of size *N* at the start of the epidemic (i.e., at time *t* = 0), denote the per-capita susceptible (*S*) disease transmission rate by *τ* (*I, N*), which we assume depends on both the number of infectious individuals *I*(*t*) and the total number of individuals *N*(*t*) in the population. In addition, we denote rates of progression from exposed (*E*) to infectious (*I*) and onto to removed with immunity (*V*) using the symbols *σ* and *γ* respectively (Figure 1). For generality, as depicted in Figure 1A, we include a population net recruitment function λ(*t*), where all these recruits are assumed to be susceptible. Later we generalize this in the context of a metapopulation structure and allow other disease classes to migrate. We also include per capita disease-induced and natural mortality rates *α* and *μ*, respectively, that accumulate in disease class D, as well as allow for the occurrence of a per capita immunity-waning rate *ν* (Figure 1). Some or all of this latter group of parameters may be zero, and only become nonzero as the scope of the analysis undertaken is enlarged. Also, as a starting point, all parameters are assumed to be constant, except for the transmission function *τ*, which at its most fundamental has the relatively simple “frequency-dependent” structure [20]

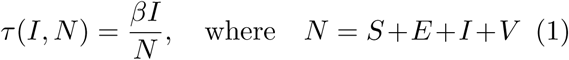

with the disease transmission parameter *β* itself constant over time. The even simpler “density-dependent” form *τ* (*I, N*) = *βI* should be used with caution, because at high population densities it is unduly unrealistic; although at low population density it can be quite useful to generalize *τ* (*I, N*) as being approximately density-dependent (e.g. see [21] and the Discussion section below).

With the above notation, the basic continuous time differential equation formulation of an SEIR epidemic process (i.e., and SEIV + D) in a homogeneous population takes the form:

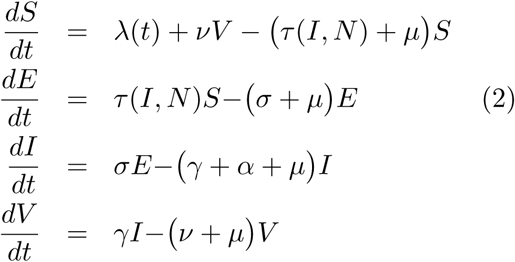

To complete the description, we need to include the relationship

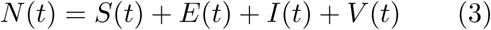

We may also want to evaluate the accumulated number of deaths *D^μ^* and *D^α^*, due respectively to natural and disease-induced causes using the equations

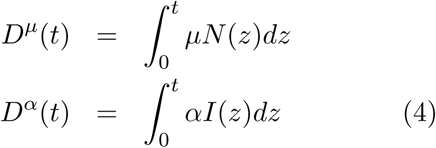

The properties of Equations 2 have been extensively studied over the past three to four decades [10–12, 22], with the most important results pertaining to the pathogen-invasion or disease-outbreak condition. This is informally derived here for the case *τ* = *βI*/*N*, λ(*t*) = *μN* (i.e., individual birth and death rates are the same) and *ν* = 0 from the following considerations. Each infectious individual infects susceptible individuals at a rate *βS*/*N* (≈ *β* when *S* ≈ *N*) over an infectious period that lasts on average for a time 1/(*γ* + *α* + *μ*). However, only a proportion *σ*/(*σ* + *μ*) of infected individuals become infectious, due to natural and disease-induced mortality rates while in state E. Thus the number *R*_0_ of susceptible individuals that each infectious individual is expected to infect at the onset of an epidemic is given by

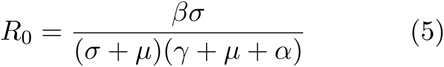

This derivation for the SEIR model, in the context of density-dependent transmission, can be found in [22]: it uses the the so-called “next generation matrix” method [23–25] to compute this result. Because an outbreak cannot occur unless *R*_0_ > 1, Equation 5 in turn implies that

Outbreak threshold:

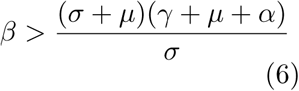

### 2.2 Numerus SEIR Continuous-Time Implementation

As an introduction to using Numerus Model Builder to code dynamical systems models, we begin with the very simple population growth model (also known as the logistic model)

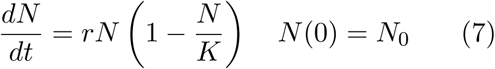

This equation can be thought of as a special case of Equations 1 and 2 when *S* (0) = *N*_0_, *E*(0) = *I*(0) = *R*(0) = 0, and λ(*t*) = *rN*(*t*) (1 − *N*(*t*)/*K*): under these conditions, *E*(*t*) and *I*(*t*) remain zero and *S*(*t*) ≡ *N* (*t*) for all *t* ≥ 0.

In Video 1 at the supporting website, the reader can find a complete construction of the logistic Equation 7 using Numerus Model Builder, with solutions generated for various values of *r* when *K* = 1. We note that there is no loss of generality in setting *K* = 1 because it can be seen to be a scaling constant that varies with the measurement units selected for *N*. (Verify this by making the transformation *x* = *N*/*K* in Equation 7.)

Using Numerus Model Builder to code up Equations 2-4, we obtain the model illustrated in Figure 2. We then ran the model for the case λ(*t*) = *μN* and *τ* = *βI*/*N* to explore the effect of *β* on solutions to Equation 2 as it increases from a value below the outbreak threshold to one above the outbreak threshold, as embodied in Inequality 6. In particular, for the parameters listed in the caption to Figure 3, it follows from Equation 5 that the threshold is

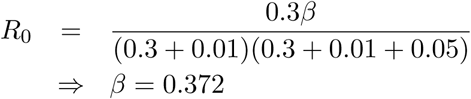

and the impact of *β* increasing from 0.37 to 0.38 is illustrated in Figure 3A. In this panel, we see that after an initial drop related to the fact that individuals entering E must first transition to I before they can begin to infect individuals in S, the solution is declining for *β* = 0.37, but ultimately growing for *β* = 0.38. In addition, as illustrated in Figure 3B, when *ν* > 0 the effect of recycling individuals from the R class back into the S class, results a slightly higher peak epidemic level that does not drop down to nearly zero before rebounding for a small echo of the first peak.

**Figure 2:**
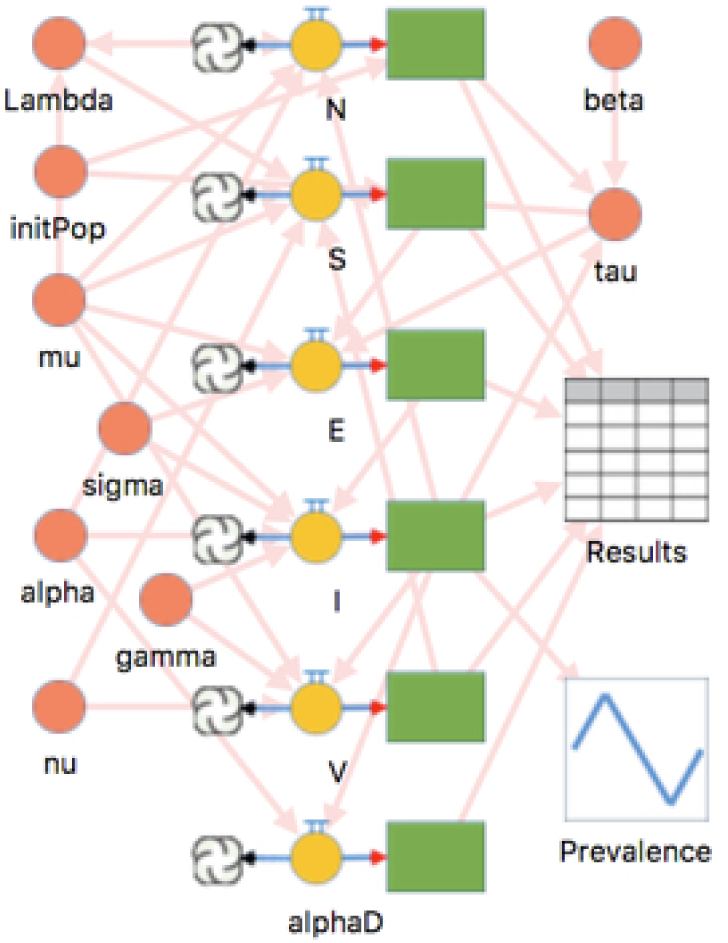
A Numerus Model Builder representation of Equations 2 to4, showing the dynamic variables as green boxes with orange circle icons that represent the differential equations of the system, and pink circles to represent input parameters or terms such as λ = *μN* or *τ* = *βI*/*N*. The grid represents an output table, and the blue in-square zig-zag a graphing tool. See Video 2 at the supporting website for more details.

**Figure 3:**
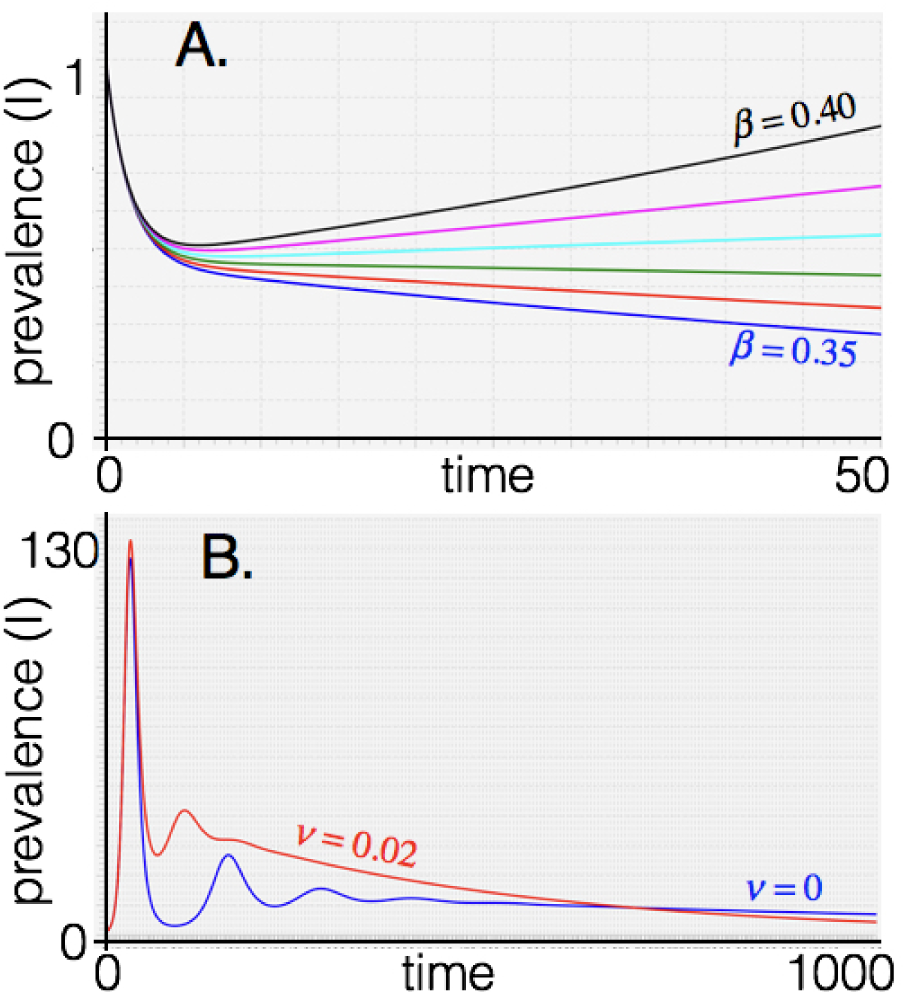
The incidence *I*(*t*) is plotted over time (A. *t* ∈ [0,50], B. *t* ∈ [0,1000]) using the Numerus model, depicted in Figure 2, to generate numerical solutions to Equations 2 under initial conditions *S*(0) = 999, *E*(0) = 0, *I*(0) = 1, and *V* = 0 for the case *μ* = 0.01, *α* = 0.05, *σ* = *γ* = 0.3, under the assumption that λ(*t*) = *μN*(*t*). In addition: in A. *ν* = 0 and *β* varies from 0.35 to 0.40 (in steps of 0.01); and in B. *β* =1 and *ν* = 0.02 (red) and *ν* = 0 (blue). See Video 3 at the supporting website for details on making sets of batch using Numerus Model Builder.

### 2.3 Discrete Deterministic Models

Discrete-time models, as represented by systems of difference equations, are computationally more efficient than continuous-time differential equation models, such as Equations 2, because discrete time models do not require numerically intensive integration. Furthermore, discrete models synchronize directly with periodically collected data: which may be daily or weekly incidence rates in fast moving epidemics, such as influenza, SARS, or Ebola; or monthly or annual rates in slower moving epidemics such as HIV or TB. In long running epidemics that have a seasonal component, such as TB [26], the effects of seasonality can only be estimated from a model if incidence rates are reported monthly or, at least, quarterly.

Discrete models present an event sequencing conundrum. For example, consider outflow from the infectious class over time interval (*t*, *t* + 1], as modeled in the third equation in Equations 2. If there are *I*(*t*) individuals in class I at time *t* then, assuming no inflow, the total number of individuals still in class I at time *t* + 1 is obtained by integrating the equation

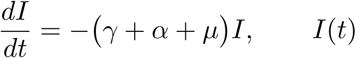

specified over (*t*, *t*+1], i.e., over one unit of time, to obtain

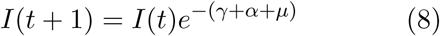

Hence the proportion of individuals that leave the infectious class over time (*t*, *t* + 1] due to recovery at rate *γ*, dying from disease at a rate *α*, and dying from natural causes at a rate *μ* is

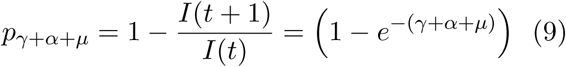

The simplest way to allocate the proportions *p*_*γ|α*+*μ*_, *p*_*α|γ*+*μ*_, and *p*_*μ|γ*+*α*_ of individuals leaving class I into those that respectively recover, die from disease, and die from natural causes is in proportion to the rates *γ*, *α* and *μ* themselves. This produces that so-called “competing rates” formulation [27]:

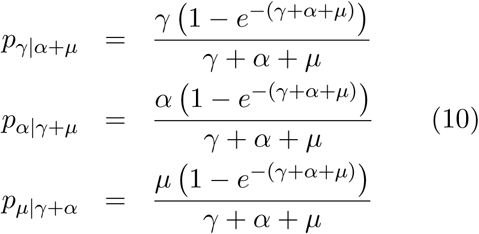

From this it easily follows that

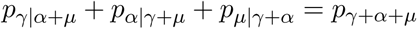

Using this notation in the context of the appropriate rates for each equation and making the assumption that

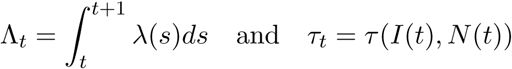

are constants that apply over time interval (*t*, *t* +1], a discrete equivalent of Equations 2 takes the form

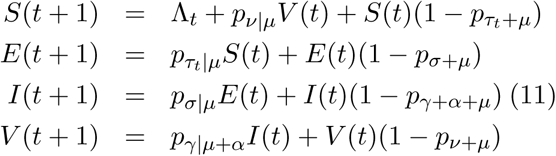

We note that solutions to this discrete system will differ from solutions to the continuous system, even for constant Λ(*t*), because *τ*(*t*) does not remain constant over the interval (*t*, *t* + 1] in the continuous model (Figure 4). However, there is no *a priori* reason to favor a continuous over a discrete time formulation because the data represent averages of discrete events (that is, transitions of individuals from one disease class to another, or the occurrence of deaths) rather than continuous, smooth flows. In fact, stochastic approaches are needed to capture the full richness of epidemiological dynamics [8,9,17] and, as discussed below, it is far easier to make discrete-time models stochastic than continuoustime models.

**Figure 4:**
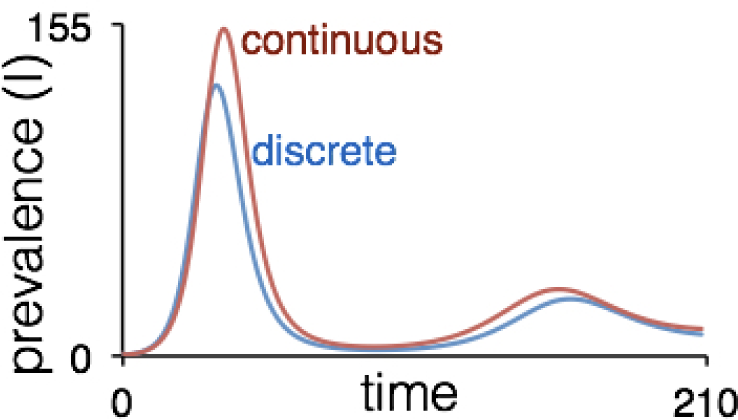
The incidence *I*(*t*) is plotted over time *t* ∈ [0,210] for the continuous (red) and discrete (blue) Numerus Model Builder coding of SEIV models represented by Equations 2 and 12 respectively for the case *τ*(*t*) = *βS*(*t*)/*N*(*t*) and λ(*t*) = *μN*(*t*) using the parameter values *β* = 1, *α* = 0.05, *σ* = *γ* = 0.3, and *ν* = 0. In the discrete model we note that Λ_*t*_ = *d_μ_*(*t*) and *τ*(*t*) is assumed constant over [*t*, *t*+1). The numerical solutions depicted here correspond to initial conditions *S*(0) = 999, *E*(0) = 0, *I*(0) = 1, and *V* = 0. See Video 4 at the supporting website for additional details on building the discrete model using Numerus Model Builder.

**Figure 5:**
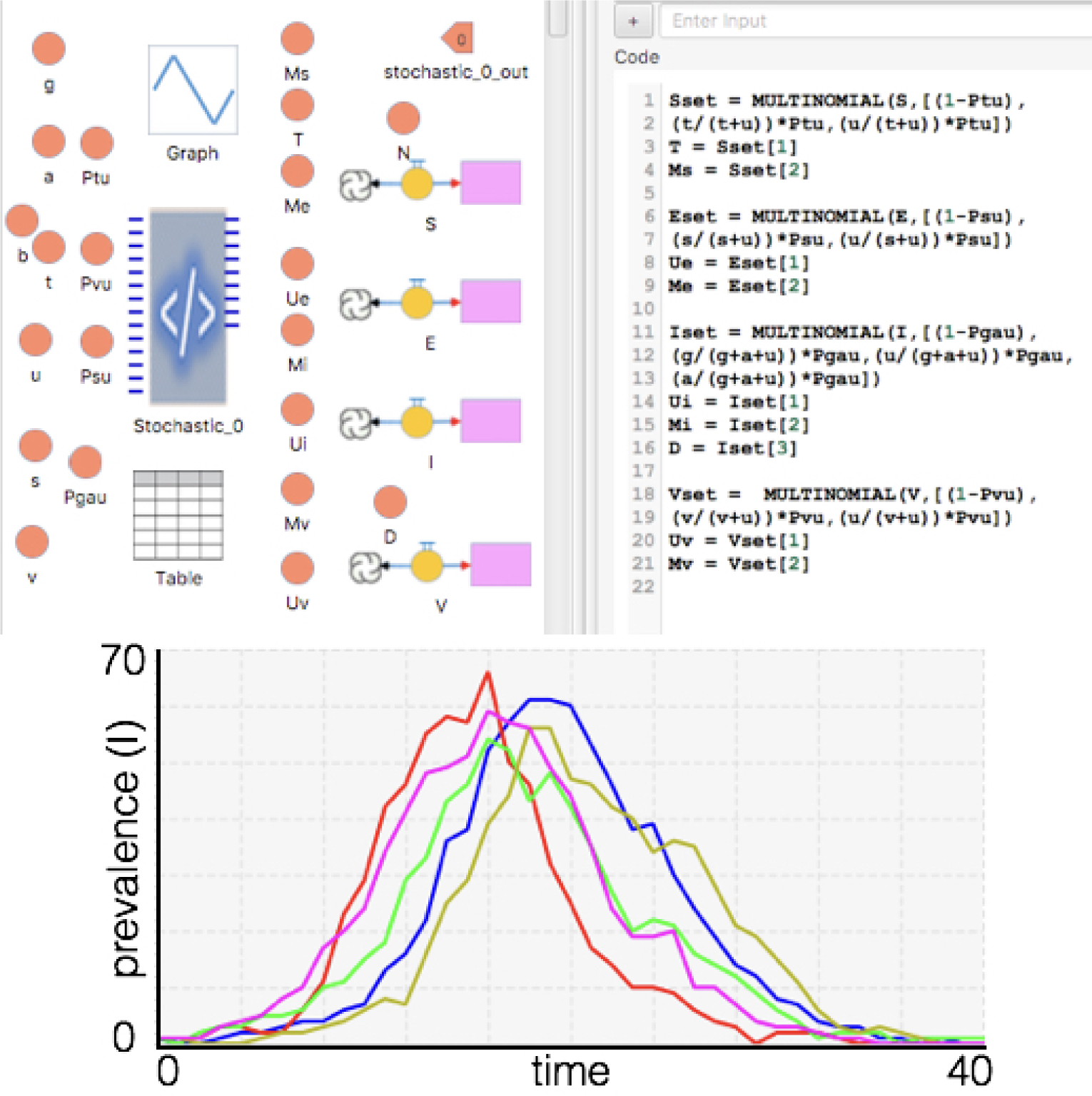
The top left panel provides a Numerus Model Builder representation of the discrete stochastic model with the stochastic chip (Stochastic_0) at its center. This chip contains the code (right top panel) for generating the multinomial distributions seen in Equations 14. The bottom panel illustrates plots of incidence from five repeated runs of the stochastic model using the identical set of parameters in each run. See Video 5 at the supporting website for more details.

To complete this discrete model, we have the discrete analogues of the continuous live and dead population variables:

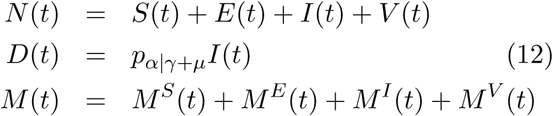

Where

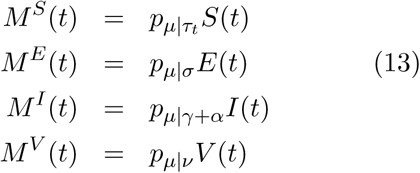

### 2.4 Discrete Stochastic SEIR Models

Before presenting a stochastic formulation of Equations 11-13, it is worth noting that one can simulate continuous systems models, such as Equations 1-4, as a stochastic process of randomly occurring events using Gillespie’s algorithm [28,29] and its refinements [30–32]. This general, event-oriented approach, however, involves considerably more computations invoking numerical integration schemes than working directly with discrete models. Further, as we stressed earlier on, a continuous-time model is theoretically no more privileged than its analogous discretized formulation that has its iteration interval synchronized with the frequency at which the data are collected.

In developing a stochastic formulation we use the notation

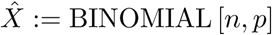

to denote that 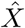 is one drawing of a binomial variable representing the number of times one of two outcomes occurs in *n* trails, when the probability of this outcome occurring in a single trial is *p* (i.e., a Bernoulli process with probability *p*). More generally, we use the notation 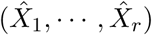 to denote one instance or one particular drawing of (*x*_1_,…, *x_r_*) ~ MULTINOMIAL [*n*; *p*_1_,…, *p_r_*], where 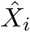 is the number of times one of *r* possible outcomes occurs over *n* trials, each have probability *p_i_* (*i* = 1,…, *r*) of occurring in any one trial.

With this notation, we can write down equations for the stochastic equivalent of the discrete deterministic model represented by Equations 11. We use the additional notation 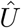 with appropriate designator subscripts to denote the number of individuals transferring between disease classes:

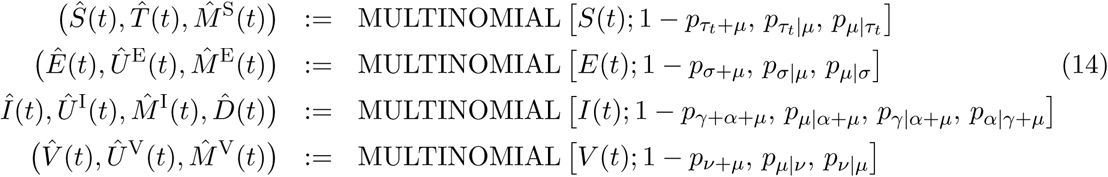

The stochastic version of our discrete SEIV model is thus represented by the following equations:

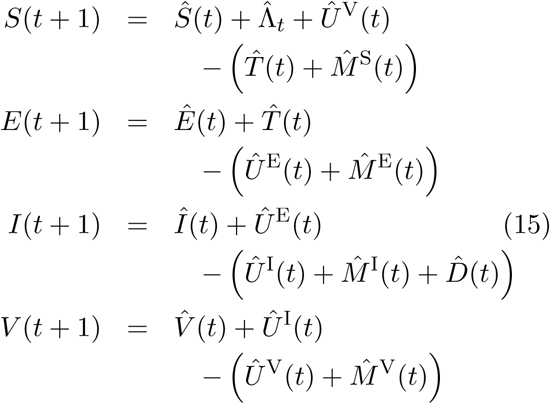

where 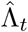 are generated from an appropriate discrete distribution, such as a Poisson distribution with expected value Λ_*t*_ determined from local population birth rates or other relevant recruitment processes.

To complete this stochastic discrete model represented by Equations 14-15, we have

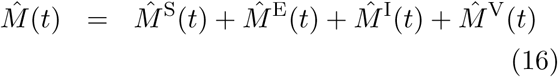

as well as *N*(*t*) in Equations 12, which is needed to calculate *τ_t_* = *τ* (*I*(*t*),*N*(*t*)) using an appropriate expression, such as given in Equation 1.

### 2.5 Weakness of the SEIR formulation

Real epidemics are far more complicated than the idealized epidemics encapsulated in the above SEIR models. Among assumptions in SEIR models used to keep them relatively simple are:

1. *The assumption of host homogeneity*. This assumptions is tenuous at best: the host’s age [33], sex [34], genetic makeup (particularly MHC locus genes) [35], physiological state [36], and history of exposure to the current and related pathogens (the latter due to cross-immunity issues [37]) all play a role in affecting the vulnerability of the host to infection, the length of time the host is infectious, and the risk of the host dying from disease.
2. *The assumption of spatial homogeneity*. This is related to the assumption that hosts contact one another at random. Contact is never random. At best, contact can be assumed to be locally random. This implies that the probability individuals contact one another over some future period is inversely related to their current distance from one another. One way around this assumption is to extend SEIR models to a metapopulation setting in which subpopulations are regarded as homogeneous and rates of exchange of individuals among subpopulations is some inverse function of the distance among the centers of these subpopulations, as discussed into Section 3 below.
3. *The assumption that the transmission rate per susceptible individual is a relatively simple function of host population and infectious class densities (or numbers)*. For example, Equation 1 assumes that total transmission has a frequency dependent form. A more general case that assumes transmission is essentially i) density dependent when population density *N* is small relative to some function-location parameter *L*, and ii) frequency dependent when the population is much larger than *L*, takes the form

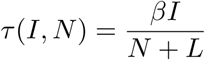 More complicated functions have been proposed [12,21], including a negative binomial expression that accounts for susceptible host aggregation [38] or the phenomenon of superspreaders [39], however, they are also limited in adhering to the following assumption.
4. *The assumption that the system is memory-less*. This assumption implies that formulations do not distinguish among individuals that have essentially spent several time periods in a particular disease state E, I or V compared to those that have just entered one of these states. This assumption can be obviated by using a discrete time models that tracks the number of days each individual has been in a particular disease state, as seen for example in a model of the 2003 Asian outbreak of SARS [40].
5. *The assumption that individuals exit disease states following exponential (continuous time) or geometric (discrete time) distributions*. We see this clearly in Equation 8 in the context of infectious individuals in the continuous time model over a single time period, which leads to the geometric rate of decay when applied iteratively over several time periods. This rather severe assumption (which implies that the highest exit proportions occur closest to entry into the disease class—or put another way, the mode of the exit distribution is at the point of entry into the given disease class) can be obviated using a box-car model or distributed-delay approach. In these formulations, infected individuals pass through disease class E by passing through a sequence of disease subclasses E_1_, E_2_,…, E_*r*_, before passing into the disease subclass sequence I_1_, I_2_,…, I_*k*_, and then finally into disease class V [41–44]. In this case, the exit distributions from E and I are no longer exponential, but are now Erlang (i.e., a subclass of the Gamma distribution). Further, the mode of the Erlang distribution becomes increasingly peaked and approaches the mean as the number of subclasses increases.
6. *The assumption that the transmission rate parameter is time independent*. Primary reasons why epidemics subside are that either the proportion of susceptible individuals in the population is reduced to the point where the epidemic can no longer be sustained (so-called threshold effect [20, 45]) or the rate at which susceptible individuals contact infectious individual during the course of an epidemic, as in the recent Ebola outbreak in West Africa [17,46], precipitously falls due to behavioral reasons as the epidemic proceeds. One approach is to assume that β has the exponential form *β*(*t*) = *β*_0_*e*^−*εt*^ (e.g. as in [47]). This is a little extreme because we should not expect *β* not to start to decline precipitously at the start of the epidemic, but only part way into the epidemic, once public awareness of the full potential of the epidemic has become apparent. In this case, a mirror-image, s-shaped curve of the form

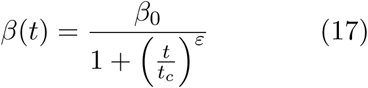

for parameters *t_c_* > 0 and *ε* > 1 is more appropriate (in a manner analogous to the onset of density-dependence, as discussed in [48]).
7. *The assumption that pathogen dose can be ignored*. The human immune system is extremely complex and takes a variable amount of time to gear up once invaded by a replicating army of pathogens, as the gear-up time depends on the condition of the host, host genetics, and prior host experience with the same and other pathogens. Small pathogen armies (i.e., low doses) are more easily contained by the hosts immune system—that is, before they can replicate to reach levels that may overwhelm and kill the host—than high doses or repeated exposure to lower doses over a short window of time. Such host-immune-system/pathogen dynamics can only be understood using models that are often more complicated than the SEIR model itself [49,50]. Further, ignoring both single and repeated dose effects may severely compromise the reliability and transferability of SEIR models fitted to one population and then applied to another population or even to the same population at a later date.
8. *The assumption that infectious individuals are equally hazardous*. Although this falls within the ambit of Assumption 1, it is worth pointing out that the phenomenon of superspreaders is well-known and that in some epidemics fewer than twenty of infected individuals may be responsible for more than 80% of transmission events [39].

## 3 Metapopulation Formulation

The first step in extending homogeneous SEIR models to a metapopulation setting is to prepare the homogeneous models by embedding them in a background population through the addition of migration processes (Figure 6A). For example, for the *i^th^* subpopulation we can add local per-capita emigration rates 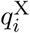 to Equations 2 for X=S, E, I, and R, to account for individuals that move in and out of the four different age classes. If we do this for all subpopulations using a set of subpopulation emigration rates 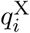, *i* = 1,…, *m*, we can then also generate a set of immigration rates 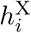 that conserves total movement numbers within the metapopulation. In this case, we need to define a movement matrix with elements 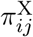 such that

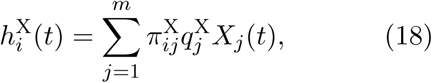

where 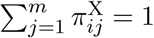 for *i* = 1,…, *m* and X=S, E, I, and V. We can then add this migration process to Equations 2, applied to the *i^th^* subpopulation in the metapopulation network, to obtain:

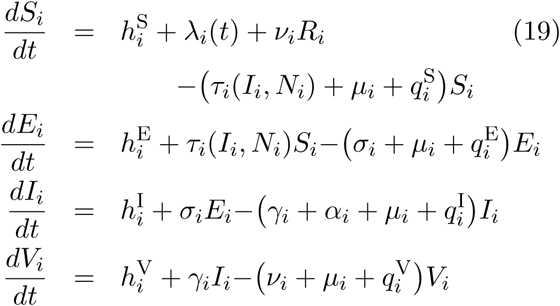

**Figure 6:**
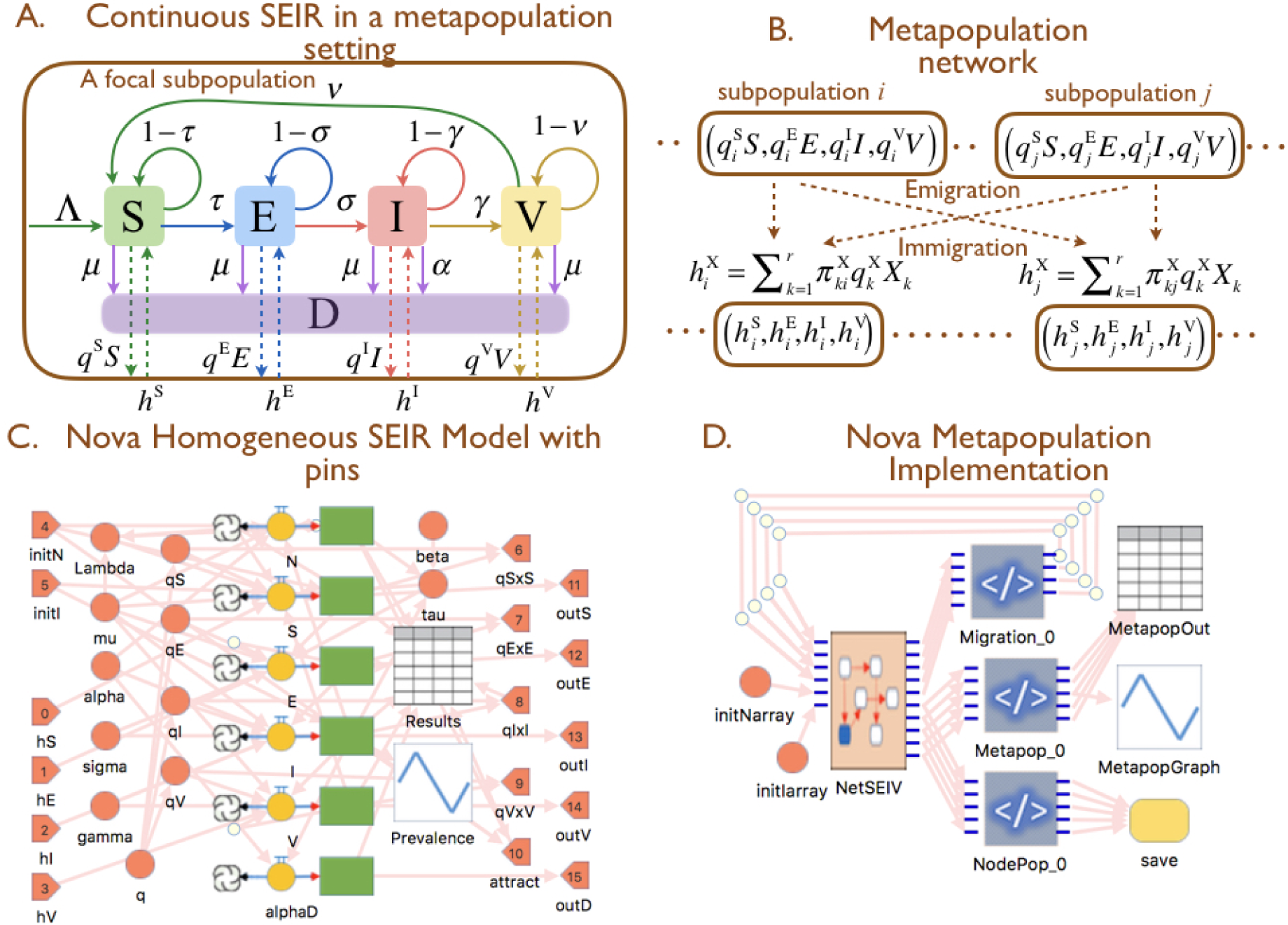
**A.** A homogeneous subpopulation with input and output flows of individuals to and from other subpopulations in the metapopulation (cf. Figure 1A. **B.** The flow network quantities and governing equation for the metapopulation as a whole. **C.** The Numerus Model Builder submodel of the subpopulation processes depicted in A, which is basically the Numerus Model Buildermodel illustrated in Figure 2 with input and output pins added as described in Video 6 at the supporting website. **D.** The Numerus Model Builder metapopulation model formulated one hierarchical level above the subpopulation model depicted in C, with the use of the NetSEIV, Migration and Metapop codechips explained in Video 7 at the supporting website. The NodePop codechip connected to the yellow “save” event container allows the one-time event of saving trajectories of all variables from all nodes at the end of the simulation.

We can further assume that the movement elements *π_jl_*(*t*) are derived from a set of connectivity strengths *K_jl_*(*t*) that reflect that relative ease with which an individual in subpopulation *j* can move to subpopulation *l* over the time interval [*t*, *t* + 1) and a set of relative attractivity values *a_l_*(*t*) that are characteristics of the nodes *l*. These attractivity values *a_l_*(*t*) are assumed to bias the movement of any individual leaving subpopulation *j* to move to subpopulation *l* with probabilities computed using the formula:

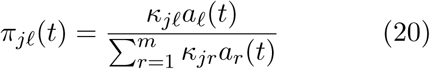

We further note that we allow *K_jl_* ≠ *K_lj_* to hold in general, though if *K_jl_* are constructed using a symmetric distance matrix with elements *ε_jl_* such that

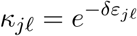

for some scaling constant *δ* > 0, then the relation *K_lj_* = *K_jl_* will hold.

The attractivity factor *a_l_*(*t*) could reflect several different aspects of the subpopulations, including their size, proportion of infected or immune individuals in the subpopulation, and so on. We will assume that two factors play a central role in determining the relative attractivity of each subpopulation: a characteristic size parameter 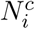 and the ratio of infectious individuals *I_i_*(*t*)/*N_i_*(*t*) for the *i^th^* population, *i* = 1,…, *m*. For example, we might assume attractivity falls off linearly from 1 to 0 with the ratio *I_i_*/*N_i_* (i.e., use the factor (1 – *I_i_*/*N_i_*)). Similarly, we might assume that the attractivity falls off as *N*(*t*) ∈ [0, ∞) varies on either side of 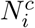 (e.g. a factor of the form 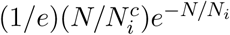 which ranges between 0 and 1 and back to 0 as N increases from 0 to 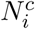 and then beyond to infinity). In this case we may define

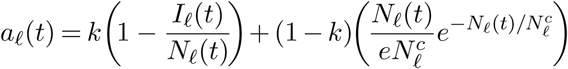

where *k* ∈ [0,1] switches the emphasis from the population size factor to the prevalence factor as *k* increases in value from 0 to 1.

The inputs 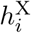(*t*) and per-capita flow rate outputs 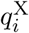 (X=S, E, I and V) for the focal *i^th^* subpopulation, can either be 0, constants, or generated using probability distributions in stochastic versions of the model. The inputs will, of course, depend on the density or number of individuals available in the environment surrounding the focal subpopulation *i*, with population structure taken into account using network or nearest neighbor concepts. In the context of discrete deterministic or stochastic models, we need to account for the per-capita flow rate outputs *q*^X^ in our competing rates formulations to obtain the extended probability for the case of rates assumed to be constant over each interval of time (though the rates themselves can vary from one time interval to the next). In this case, we augment the proportions/probabilities in Equation 10 to define the following terms for constructing the infectious class equation in the where *i^th^* subpopulation

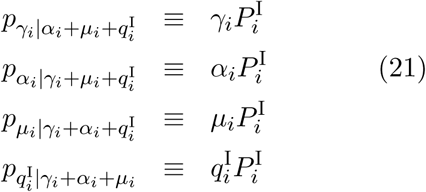

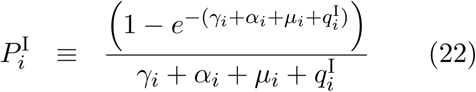

and, as before, it follows that

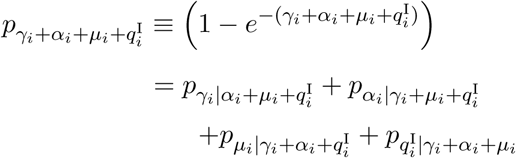

with similar expressions following for the susceptible, the exposed and the immune classes ex-pressed in terms of 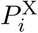, X=S, E, I, and V following the patterns of Equations 21 and 22

These expressions can be used to write down an extended version of the deterministic discrete model given by system of Equations 11 or of the stochastic model given by system of Equations 14 and 15. By way of illustration, using 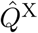 to represent the proportion of individuals leaving class X=S, E, I, and V due to immigration, Equations 14 now become (dropping the argument in *t* and the subscript *i*)

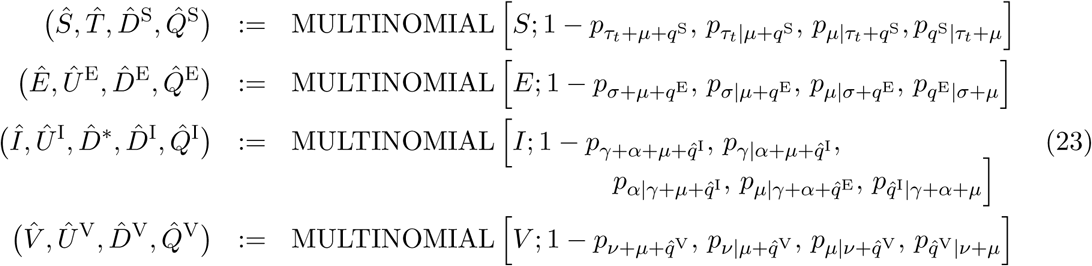

Thus, in the case of the *i^th^* subpopulation, the actual number of individuals leaving the different disease classes during the time interval [*t*, *t* + 1) (after sampling has been applied) are:

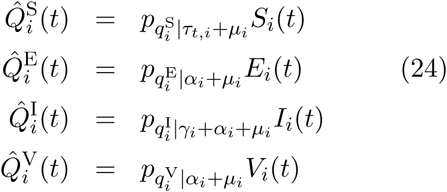

Assume the only source for individuals immigrating to a subpopulation during the interval [*t*, *t* + 1) are those emigrating from all of the other subpopulations. Given this, as in setting up Equations 18 and 20, we can now express the emigrants 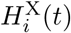 (*i* = 1,…, m) in terms of the immigrants 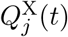 (*j* = 1,…, *m*, for each X=S, E, I, or R), and the parameters *π_jl_*:

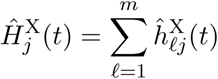

X=S, E, I, and V where the individual 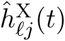 are generated from the drawings

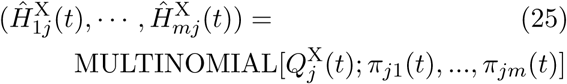

With this process completed, we then obtain the following extended version of Equations 15

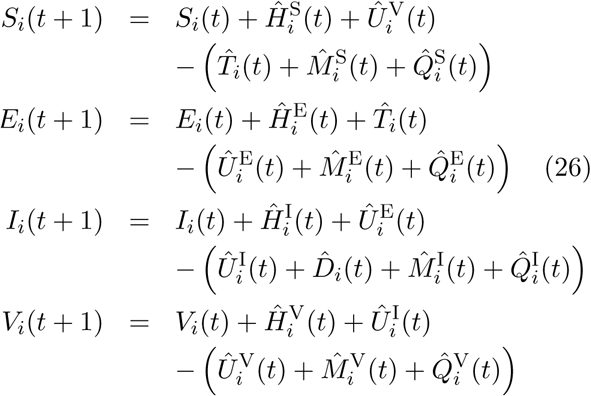

The recruitment numbers 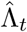, generated during each interval (*t*, *t* + 1], are drawn from an appropriate discrete stochastic process. The simplest is a Poisson process with expected value Λ_*t*_, where the latter is determined by local population birth rates or other processes generatingnew individuals. Illustrative simulations of the metapopulation model depicted in Figure 6 for the case of 6 locations strung out in a row, where individuals can only move between neighboring locations are provided in Figure 7.

**Figure 7:**
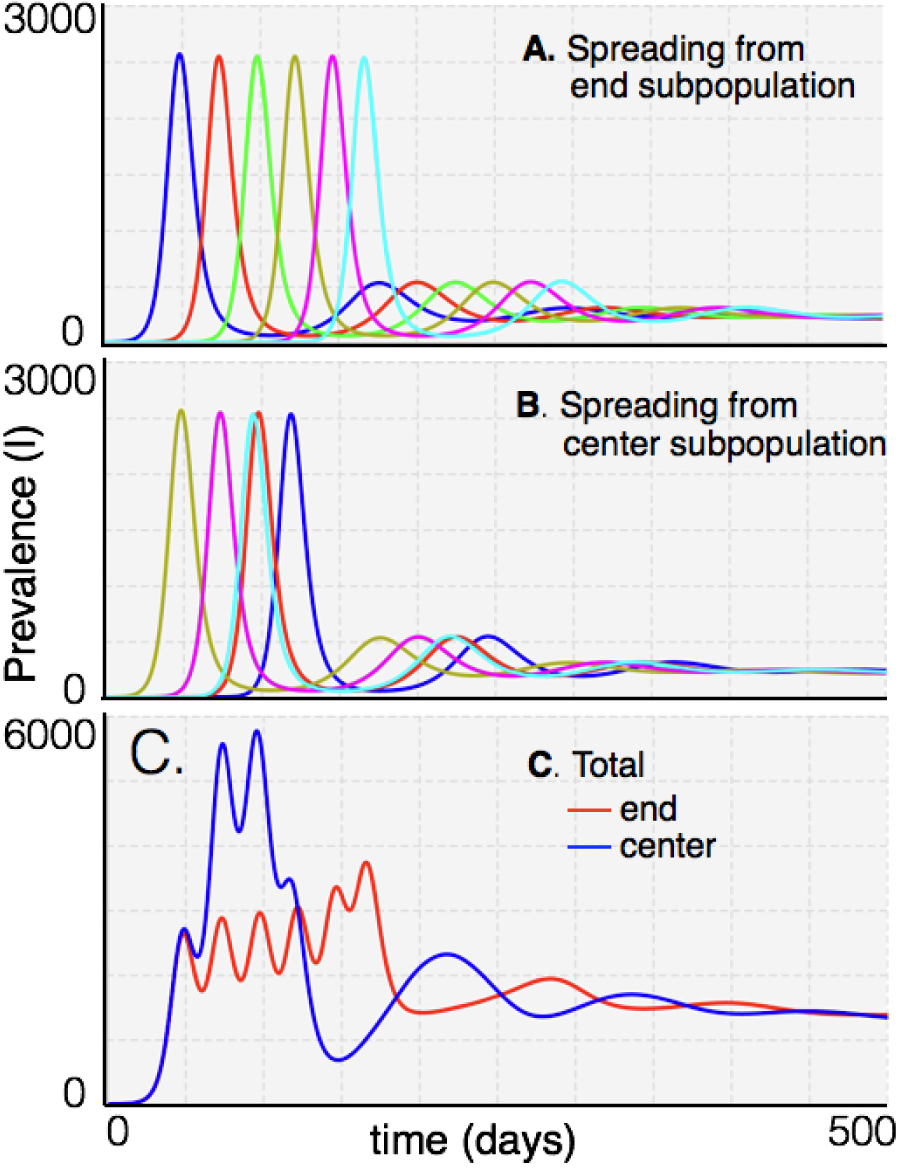
Fig: Prevalence plots predicted by a continuous-time deterministic metapopulation model when the index case starts out in one of six possible locations (each with *S*(0) = 20, 000 and other classes at 0, except for the location that has *I*(0) = 1 and *S*(0) = 19, 999), where the locations numbered from 0 to 5 are strung out in a straight line in the order 0, 1, 2, 3, 4, 5, and individuals flow only to subpopulations that are their immediate neighbors. Subpopulation prevalence rates are plotted in **A**. (index case in subpopulation 0) and **B**. (index case in subpopulation 2) with the subpopulation containing the index case clearly leading the outbreaks in closest and the next closest neighboring populations. Total prevalence is plotted in **C**. for outbreaks with two index cases that are either the end two (red plot) or center two (blue plot) subpopulations.

## 4 Fitting Models to Data

Methods for fitting epidemiological and other types of dynamical systems models to data is a vast topic that requires mathematicians, statisticians, and engineers around the world working on the problem full time. Here we only touch the surface of the topic and discuss how Numerus Model Builder can be used to address the issue under relatively straightforward and manageable situations (i.e., not too many equations and with a few parameters at most free to vary during the fitting procedure). A gentle introduction to the field of fitting population models to data is provided by Hilborn and Mangel [51]. We will avoid the issue of model selection itself [52, 53]—i.e., if one fits various models to data with different numbers of parameters, then how does one decide which of these models best fits the data in an information theoretic sense—since this is also a research topic that attracts considerable attention.

Fitting dynamic models to a set of observations **Y** = {*Y*_1_,…, *Y_n_*}, where the index *i* in *Y_i_* refers to time *t* = *i*, *i* = 1,…, *n*, typically involves generating a set of comparable values 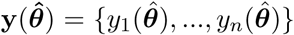 from a model that has a set of m parameters ***θ*** = {*θ*_1_,…, *θ_m_*}, where 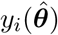 is the value of some variable in the model at time *t* = *i* when the parameter values are 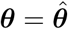.

The two dominant approaches to fitting models to data are least-squares estimation (LSE), which has been largely supplanted by the maximum likelihood estimation (MLE) [54]. The latter is typically embedded in a Markov Chain Monte Carlo (MCMC) algorithm that constructs a probability distribution for ***θ*** using Bayes theorem [55–58]. MCMC requires the likelihood function to be known. This can be obviated, though, by assuming the distribution of model outcomes to be Poisson (as we do below), using likelihood-function-free methods [59], or using approximate Bayesian approaches [60].

LSE methods involve minimizing the sum-of-squares residuals (or error) measure

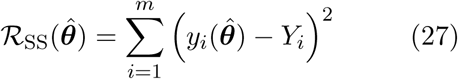

On the other hand, MLE methods that assume model outcomes are Poisson, but with a different Poisson mean 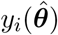 for each data point *Y_i_*, *i* = 1,…, *t*; involves maximizing the log-likelihood function

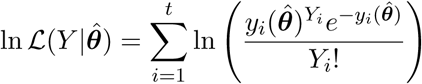

or minimizing its negative, which can be written for 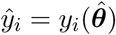 as

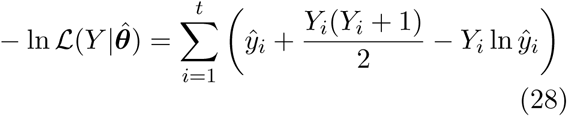

Here, for purposes of illustration, we fit our deterministic SEIV model to the Sierra Leone Ebola weekly incidence data [61] using both LSE and MLE approaches (Figure 8: cf. fits obtained in [47]). When fitting such data the appropriate initial conditions are generally uncertain because detection of the putative index case does not generally pin down the start of the epidemic: the actual index case may often go undetected and the number of individuals in class E at the time of the first case is also unknown. Thus, as part of the fitting procedure, we allow the initial values *E*(0), *I*(0) in the model to be fitted to the data. To keep the dimensions of the fitting problem down, however, we set

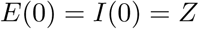

and the search for the best fitting value of *Z*. Another imponderable is the actual number of individuals *N*(0) at risk at the start of the epidemic. Thus we also treat *N*(0) = *N*_0_ to be an optimization parameter, though we set *V*(0) = 0 under the assumption that if some individuals in the population were immune to Ebola at the start of the epidemic, this would be reflected in a lower-valued estimate of *N*_0_. Thus *N*_0_ should be interpreted as the “initial population at risk” rather than actual popultion size. Also, in preliminary runs of our optimization algorithm (i.e. when fixing diffent combinations of parameters and solving reduced parameter set problems), the difference between optimal values for *σ* and *γ* under variety of settings always lead to optimal values that differ by less than a few percent. Thus to further reduce the dimension of the optimization problem, we set *σ* = *γ* during the optimization procedure. Further, we also fixed *μ* and *γ* (which in problems like this can be estimated outside of the incidence data) to 0.001 and 0.05 respectively, as well as setting λ = *ν* = 0. A more rigorous fitting of the Sierra Leone data would need these parameters to be properly estimated ahead of time, but our purpose here is to demonstrate different aspects of the fitting procedure, rather than undertaking an in depth analysis of the epidemic itself.

**Figure 8:**
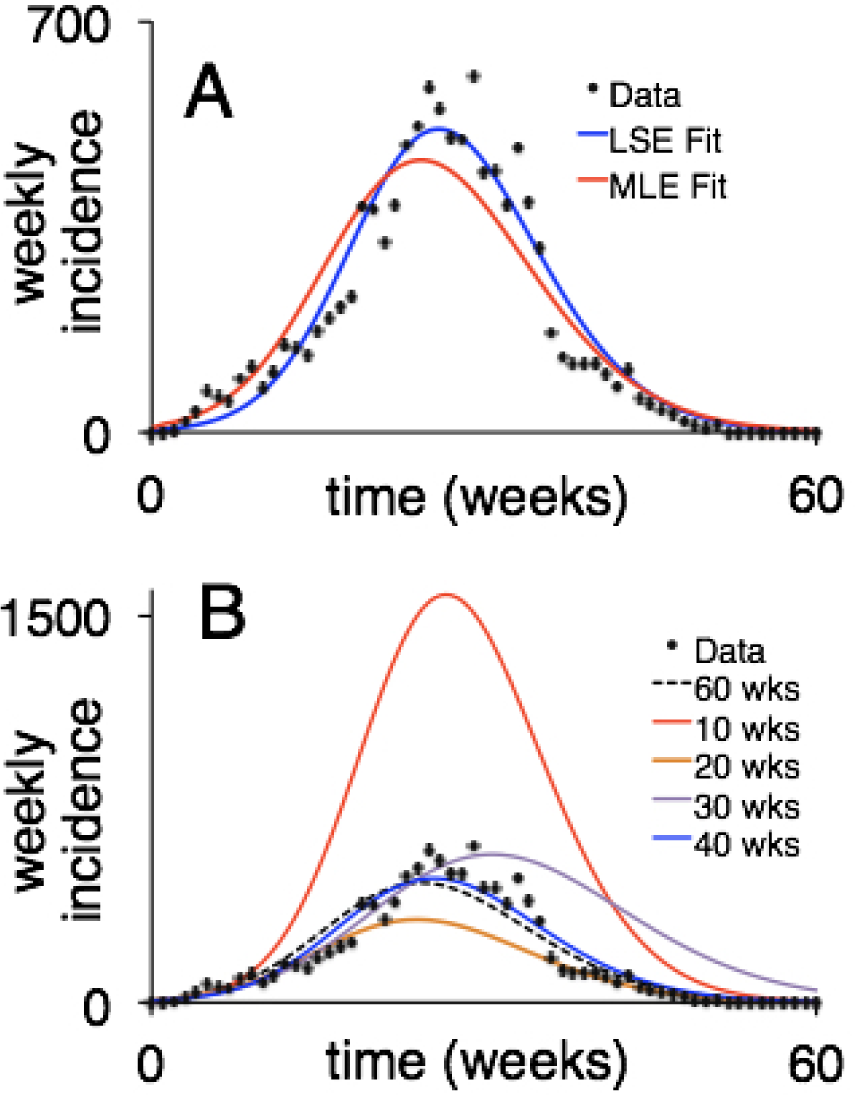
The SEIV discrete time model given by Equations 11 with fixed parameters *ν* = 0, *μ* = 0.001, *α* = 0.05 and Λ_*t*_ =0 (cf. Figure 4) has been fitted to Ebola data from the Sierra Leone 2014 outbreak in which more than 10,000 cases occurred during the course of an approximately one-year period [61]. A. The blue and red curves are the best fit LSE (cf. Equation 27 and MLE (cf. Equation 28 obtained with the optimal parameter sets given in the text. B. The black dotted line is the MLE fit, as in Panel A, with the red, orange, purple and blue plots, simulations obtained after obtaining the best MLE fits to the first 10, 20, 30 and 40 weeks of incidence respectively. See Video 9 at the supporting website for more information on how to set up and run optimizations on this model using Numerus Model Builder.

With the above parameter and model settings, the best-fitting remaining parameter values, de-noted by asterisk and obtained using a Nelder-Mead minimization algorithm, were

LSE:

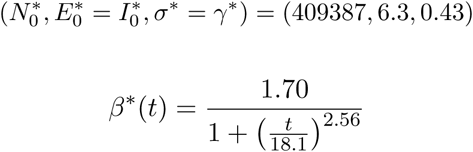

MLE:

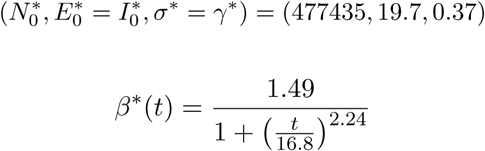

In Figure 8A we see that the LSE and MLE provide similar fits, though both provided relatively poor fits to the first third of data: these data reflect a more linear than the exponential initial phase which is uncharacteristic of homogeneous SEIR models, particular those that have no spatial structure. In general, we should not expect and SEIR model to fit the data particularly well because many of the assumptions inherent in the SEIR model, as discussed in the previous section, are violated to some often-unknown and likely-large degree.

Though the fits are similar—both, for example predict an initially *β* around 1.5 that drops to half that level in just over two weeks after the first cases have been detected—the LSE fit predicts an 14% smaller initial population at risk than the MLE fit. Thus the errors associated with this estimation can be quite large, and require a Bayesian type Markov Chain Monte Carlo (MCMC) approach to estimate them. Further, as is evident from our plots in Figure 8B, if we use the first *T* weeks of data, *T* = 10,20,30, and 40,to fit the model, we see that fitting the first 10 weeks (red curve) greatly overestimates the final size of the epidemic, while fitting the first 20 weeks somewhat underestimates the final number of cases. Before leaving this example, we note that in several runs of the optimization algorithm, different starting conditions converged to different solutions, thereby indicating that some of these solutions are local rather than global minimum. When this happened, we selected the solution that gave the lowest log-likelihood value, but our searches were not sufficiently exhaustive for us to be sure that we had found the global minimum for each case.

## 5 Discussion and Conclusion

What is the value of building SEIR models in anticipation of, during, or after an epidemic outbreak, given the level of accuracy that can be expected from such models? As with all models of complex biological systems centered around organisms, populations or communities, the answer is the same: models provide a framework for obtaining insights into dynamic population processes that could not be obtained without them. Further, they provide a means for exploring and assessing the efficacy of interventions and other types of management actions designed to protect, conserve, or exploit the populations under consideration. In the context of epidemics this has certainly been true with regard to implementing vaccination [17,62,63] and quarantine programs [64], assessing the effects of behavior [65], case detection [66], and treatment rates [67,68], managing the logistics of setting up treatment facilities during the course of epidemics, evaluating the efficacy of educational [69] and prophylactic campaigns [70], as well as drug-delivery programs that reduce the risk of producing drug-resistance pathogen strains [71].

Models are also the best tools for guiding our response to an outbreak once it has begun. Although, after 10 weeks, a fit to the Sierra Leone data set would have substantially underestimated the problem at hand, the fit at 20 weeks provided a much better ball park assessment of the final size of the Sierra Leone outbreak than could have been obtained with non dynamic modeling efforts. This remained somewhat true at 30 weeks and certainly so at 40 weeks, despite all the known serious violations of an SEIR model applied to an inhomogeneous, spatially-structured population. Thus SEIR models remain an important tool for managing epidemics, provided we treat predictions from such models with circumspection.

## Acknowledgements

The development of Nova, which is a precursor to Numerus Model Builder, was supported by NSF grant CNS-0939153 to Oberlin College (PI: RS) and NSF-EEID grant 1617982 (PI: WMG). We thank John Pataki of Logical Laboratories for his considerable help and input into creating and supporting the ‘Nova Modeler’ website (URL: http://novamodeler.com/). We thank Juliet Pulliam for hosting WMG at the South African Center for Epidemiological Modeling at the University of Stellenbosch, South Africa, during the initial drafting of the ms.

## Contributions

The initial text was drafted by WMG and the Numerus model builder platform was created by RS. WMG, RS, OM, and KT built the models. WMG created the figures, HSY and KT created the videos. All authors read and edited the manuscript.

## List of supporting online material

To enable the reader to run all our models, we have provided the 6 models we used to generate the results presented in our paper: they are available at https://www.dropbox.com/sh/i5bh700pbfijrcq/AAB2HBEs7MA4ir4CRJyDhb4Ia?dl=0. These models are listed below together with the videos (which can be downloaded at YouTube using the url links provided) that explain some of the details on how the models where built using the Numerus Model Builder (NMB) Platform. A free Mac version of NMB Light is available at https://www.dropbox.com/s/6kyli8zqz0yl8vs/NumerusMBL-0.056.dmg?dl=0, and a free Windows version of NMB Light is available at https://www.dropbox.com/s/wbn06fvubt9ovs7/NumerusMBL.056_setup.exe?dl=0. NMB allows users to build their own models or to modify the structure of the models listed below. It also allows users to run the models, input data as a comma separated string (Model 6, see Video 9 for details) and set different parameter values in each run using sliders.

### Video 1 and Model 1

Building and running a logistic differential equation model. Model used is model1_logistic.nmd. Video link is: https://www.youtube.com/watch?v=bUTvWWqVzUY.

### Video 2 and Model 2

Building and running a continuous time SEIR model. Model used is model2_SEIR_cont.nmd. Video link is: https://www.youtube.com/watch?v=v6EIvPrE5Dk.

### Video 3 and Model 2

Demonstrating how to make batch runs to generate the simulations plotted in Figure 3A. Model used is model2_SEIR_cont.nmd. Video link is: https://www.youtube.com/watch?v=-YywvTp_scs.

### Video 4 and Model 3

Setting up a discrete SEIV model. Model used is model3_SEIR_disc.nmd. Video link is: Video link is: https://www.youtube.com/watch?v=_wpHNl_acEQ.

### Video 5 and Model 4

Setting up a stochastic SEIV model. Model used is model4_SEIR_stoch.nmd. Video link is: https://www.youtube.com/watch?v=VAGtIvq_YCo

### Video 6 and Model 5

Adding pins to the continuous SEIV model and implement model as a chip at a higher hierarchical modeling level. Model used is model5 _SEIR_metapop_cont.nmd. Video link is: https://www.youtube.com/watch?v=bZLH8HAF2LQ.

### Video 7 and Model 5

Using the SEIV model in Video 6 as a submodel in a metapopulation network setting. Model used is model5_SEIR_metapop_cont.nmd. Video link is: https://www.youtube.com/watch?v=k8IEfIQCM9o.

### Video 8 and Model 5

Setting up a net-work model using the Numerus Model Builder mapping tool. Model used is model5_SEIR_metapop_cont.nmd. Video link is: https://www.youtube.com/watch?v=CIvoDM6a-HE.

### Video 9 and Model 6

Demonstrating how to use the least-square fitting tools and finding the parameter *β* in the SEIV discrete model the best fits the Sierra Leone 2014 Ebola incidence data. Model used is model6_SEIR_disc_datafit.nmd. Video link is: https://www.youtube.com/watch?v=7gLMpzb-R7Q.

